# G-ALN: a GPU-accelerated BWA-aln algorithm for palaeogenomics

**DOI:** 10.64898/2026.10.18.759956

**Authors:** Bilal Sharif, Krister Hedfors, Tom van der Valk, Peter D. Heintzman, Love Dalén

## Abstract

**Summary:** The most commonly used method to achieve optimal mapping for short and damaged ancient DNA (aDNA) reads is the BWA-aln algorithm with seeding disabled. However, this CPU-based algorithm is computationally demanding, as the aligner must tolerate mismatches and indels across the entire read. To improve alignment speed, we present G-ALN: a GPU implementation of the BWA-aln and indexing algorithms. G-ALN first searches all reads on GPU cores, with reads that need a long search being shared dynamically between the GPU and any spare CPU cores. Every read is searched using the same algorithm as BWA, so the output is byte-identical, which we confirmed here using woolly mammoth and Neanderthal datasets with aDNA-specific parameters. In our benchmarking, G-ALN indexed a reference genome ~32 times faster than BWA, and read alignment speed increased ~6-fold. G-ALN was developed through AI-assisted coding (Claude Opus 5.5). It processes thousands of reads in parallel on each GPU and aligns more than twice as many aDNA reads per unit of energy. These results show that G-ALN offers a practical route to using GPU resources in established aDNA analysis workflows.

**Availability and Implementation:** G-ALN is implemented in C++ and CUDA and is freely available under the GPLv3 license at https://github.com/bilalbioinfo/galn. It requires an NVIDIA GPU with compute capability 7.0 or higher and CUDA 11.8 or later.

**Supplementary information:** Supplementary data are available online. The sequencing read datasets used for benchmarking are available at Zenodo repository: https://doi.org/10.5281/zenodo.23455049.

## 1. Introduction

High-throughput DNA sequencing has transformed genomics research by making it possible to generate ever-increasingly large amounts of sequence data rapidly and at low cost. As a result, this has exacerbated the crucial and computationally demanding step of read mapping, *i.e*. the alignment of short DNA reads to a reference genome (Robinson, Harkin, and Shukla 2021, Sharif *et al*. 2026). These computational demands are especially high for aDNA datasets, as the short, fragmented nature of aDNA (typically below ~80 bp) means that more reads are needed to reach a given genomic coverage as compared to longer reads derived from modern DNA. Moreover, post-mortem DNA damage, as well as evolutionary divergence from modern reference genomes, creates higher levels of mismatches compared to modern DNA data. Insertions and deletions can further complicate alignment by requiring that alignment tools consider gapped solutions. Ancient DNA sequencing libraries also contain substantial non-endogenous DNA from microbes and other sources, adding reads that are considered for alignment but are without a match to the target reference genome (Schubert *et al*. 2012, Oliva *et al*. 2021, Dolenz *et al*. 2024).

BWA-aln, also known as BWA-backtrack (Li and Durbin 2009), is the most widely used aligner for aDNA population genomic studies. Commonly used aDNA settings disable seeding and allow for additional mismatches and gaps (Green *et al*. 2010, Schubert *et al*. 2012). BWA-aln is also incorporated into most established aDNA workflows, including DNAharvester, EAGER and PALEOMIX (Schubert *et al*. 2014, Fellows Yates *et al*. 2021, Sharif *et al*. 2026). Benchmarking has shown that BWA-aln performs better for aDNA compared to other widely used aligners, achieving higher alignment precision and lower reference bias, especially for ultra-short aDNA reads of ~35-50 bp (Oliva *et al*. 2021, Dolenz *et al*. 2024).

However, the performance of BWA-aln comes at a high computational cost. BWA-aln uses a best-first backtracking search, and disabling seeding increases the search needs for each read, since the aligner needs to handle mismatches and indels across the full length of the read (Li and Durbin 2009). The resulting workload can vary greatly among reads, those that near-perfectly match the reference genome require relatively little search, while reads with high divergence require much more. Thus, mapping large aDNA datasets generally consumes substantial compute time.

A non-trivial limitation is that BWA-aln runs exclusively on Central Processing Units (CPUs). At the same time, AI-driven investments are rapidly expanding Graphics Processing Units (GPU) capacity in high-performance computing (HPC) systems. This creates a growing mismatch between the increasingly available hardware at HPC clusters and the hardware that BWA-aln uses, making access to sufficient CPU capacity a key constraint on scaling aDNA analyses.

Graphics processing units have the potential to increase mapping efficiency through the processing of many independent reads on thousands of parallel cores. Although BWA-mem has recently been ported to GPU, providing substantially higher speeds (Pham, Tu, and Lv 2023), this algorithm is poorly suited to mapping ultra-short aDNA reads (Dolenz *et al*. 2024). Moreover, an early attempt to exploit GPU parallelism, BarraCUDA, ported the BWA-aln component to NVIDIA GPUs and reported faster alignment with comparable fidelity (Klus *et al*. 2012). However, its latest release (0.7.0, 2015) no longer compiles with current CUDA versions without code changes. Crucially, we found that BarraCUDA was unsuitable for aDNA as it overrides the user’s seed length, preventing disabling of the seed, and outputs are not identical to BWA-aln. A modern GPU implementation would therefore need to use current accelerators efficiently, handle the widely varying search effort of individual reads, and reproduce BWA-aln exactly so as to enable seamless integration into existing workflows.

Here we present G-ALN (pronounced ‘gallon’), a software that runs BWA-aln and indexing on GPUs, resulting in at least 6.3-fold and 32-fold higher speeds, respectively, compared to when run on state-of-the-art CPU nodes. We also show that G-ALN generates files that are byte-identical to those of the latest version of BWA, leaving all downstream tools unaffected.

## 2. Implementation

### 2.1 An exact reproduction of the bwa index and aln algorithms

G-ALN (*galn*) reproduces every step of the *bwa index* and *bwa aln* algorithms and produces byte-by-byte identical output files to its BWA equivalents. The *galn index* algorithm outputs the same five files as *bwa index* (using the same .pac, .ann, .amb, .bwt, and .sa file extensions). It uses the same code as BWA to convert a FASTA file into a 2-bit sequence (.pac), and then builds the Burrows-Wheeler transform (BWT). It uses the open-source libraries libcubwt on the GPU for genomes smaller than about 1 Gbp and libsais64 on the CPU cores for larger genomes (Grebnov, n.d.-a, n.d.-b). Finally, the occurrence counts and the sampled suffix array are computed on the GPU. Whereas the *bwa index* algorithm samples the suffix array with one sequential walk over the whole text, *galn index* instead starts an independent short walk at every sampled position and joins the pieces with a parallel list-ranking step that takes less than a second for a 3.4 Gbp genome.

The *galn aln* algorithm uses the same read parser and batching rule as BWA (262,144 reads per batch, where the longest read sets the search limits of all reads in a batch). The core BWA search function (bwt_match_gap) and the occurrence counting by the BWT are ported line-by-line into code that is compiled for both CPUs and GPUs, keeping the identical BWA integer truncations, unsigned comparisons, and floating-point rounding. Each read is searched end-to-end by a single engine (one GPU thread, one GPU warp of 32 threads, or one CPU thread; **Fig. 1a**), following the same order of search steps as bwa aln. Therefore, the results do not depend on whether a read was searched on the GPU or on a CPU core. The resulting SAI file, storing the alignment coordinates per read, is written in the same order as the fastq-file input order. Although not used when running the *galn aln* algorithm, we store a -t (number of threads) flag in the file header, in order to mimic the information that is stored in the file header by *bwa aln*, allowing for byte-by-byte comparisons. Users who want to confirm byte-for-byte identical output should give *galn aln* the same -t value they used for their *bwa aln* run (using *bwa* version 0.7.19).

**Figure 1.**
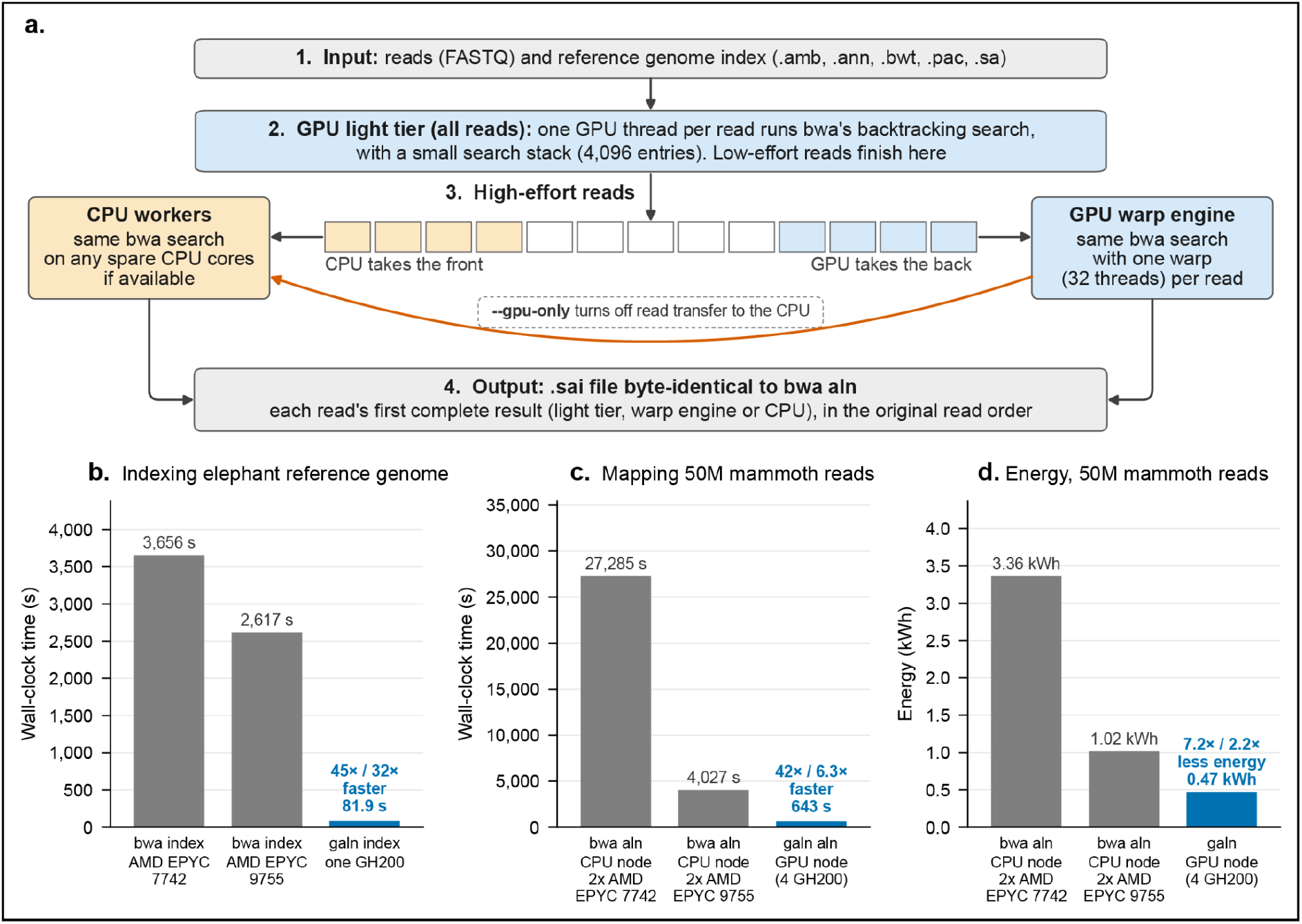
G-ALN design and performance benchmarking. (a) Overview of the *galn aln* algorithm that uses a GPU light tier to finish low-effort reads, and CPU workers and the GPU warp engine to share the high-effort reads. The output SAI files are identical to those produced by *bwa aln*. (b) Wall-clock time to index the Asian elephant reference genome (mEleMax1, 3.4 Gbp) using *bwa index* or *galn index*. (c) Wall-clock time to map 50 million reads from a woolly mammoth shotgun library (MD090; Díez-Del-Molino et al. 2023) to the Asian elephant genome with aDNA settings (-l 1024 -n 0.01 -o 2), using *bwa aln* or *galn aln*. (d) Energy used for the analyses in panel (c), from the power sensors available to users and not directly measured with an energy meter. In (b), (c), and (d), BWA was run on a CPU node with two AMD EPYC 7742 processors (128 cores; Dardel) and on a CPU node with two AMD EPYC 9755 processors (256 cores; Arrhenius), and G-ALN on a GPU node with one or four NVIDIA GH200 chips (Arrhenius).

### 2.2 Searching read alignment space using a hybrid GPU and CPU approach

G-ALN searches read alignment space in two stages due to the uneven computational effort required per read (**Fig. 1a**). First, every read is searched by a single GPU thread with a small search stack (4,096 entries) in the GPU light tier. Low-effort reads, which are aligned without exploring the whole search space and have no or few mismatches finish here. The remaining high-effort reads go into one list per GPU. A persistent GPU warp engine kernel searches one high-effort read per warp. The 32 threads of a warp run the search in lockstep, which avoids thread divergence, and split the index memory loads between them. Stack entries are packed into 16 bytes instead of the 32 bytes of BWA, so that a warp can hold more than the default queue limit for BWA of two million entries. Concurrently, if spare CPUs are available on the GPU node, CPU worker threads take high-effort reads from the other end of the same list. The GPU and the CPU exchange their positions in the list about once per millisecond with small asynchronous copies, so no shared or unified memory is required. Each CPU worker advances eight reads in turn and prefetches the index blocks they will need next, which hides much of the memory latency of the search. When the GPU list is completed, it hands its highest-effort unfinished read alignment searches to the CPU workers, and idle CPU workers also re-run reads the GPU is still considering. The first finished copy is kept for each read and, since all copies are identical, the output never depends on timing or how the job was parallelized.

## 3. Benchmarking

### 3.1 Benchmarking setup

We compared BWA 0.7.19-r1273 (Li and Durbin 2009) with G-ALN 0.2.1 on the NAISS Arrhenius and Dardel (PDC) clusters. BWA was run on full CPU nodes: two AMD EPYC 9755 (256 cores) on Arrhenius and two AMD EPYC 7742 (128 cores, 256 threads) on Dardel, with *bwa aln -t 256*. G-ALN ran on a full Arrhenius GPU node with four GH200 chips. In G-ALN, -t only sets the value written in the SAI header, so we set it to 256 to match bwa aln.

We indexed the Asian elephant (mEleMax1; 3.40 Gbp; GCF_024166365.1) and human (GRCh38.p14; 3.3 Gbp; GCF_000001405.40) reference genomes with *bwa index* on the CPU nodes stated above and with *galn index* on one GH200. We then sampled 1, 20 and 50 million reads each from the woolly mammoth MD090 (Díez-Del-Molino *et al*. 2023), the steppe mammoth Adycha (van der Valk *et al*. 2021), and 1, 2 and 5 million reads from the Neanderthal Chagyrskaya 8 (Mafessoni *et al*. 2020) sequencing data. For the sampled mammoth datasets, we merged raw reads and trimmed adapters in each dataset using fastp 1.4.0 (Chen *et al*. 2018), and retained the merged reads for alignment. For the sampled Neanderthal datasets, we extracted reads from the published BAM files. We aligned these datasets to their appropriate reference genome under both default and aDNA (*-l 1024 -n 0*.*01 -o 2*) settings (Schubert *et al*. 2012).

We measured aligner wall-clock time and energy consumption for generating the 50-million-read alignments for woolly mammoth MD090 under aDNA settings. Wall-clock time was measured with the Unix time command. We measured energy consumption from sensors users can read. On the GPU node, we integrated the power of the four GH200 modules, sampled about every 0.5 s. On the Arrhenius CPU node, we integrated the power of the two CPU sockets in the same way and added an estimate for memory. On Dardel, we read the whole-node energy counter (Cray pm_counters) before and after the run.

We tested exactness between aligners for all dataset combinations using MD5 checksums. For these tests, we ran BWA-aln on the CPU node, and G-ALN both in its default mode and with *--gpu-only*.

### 3.2 Output identity, speed and energy usage between aligners

For all dataset combinations tested, the G-ALN runs wrote output that was identical to that of its BWA equivalent. This included all index files and SAI files (**Table S1**). For the 50 million woolly mammoth reads with aDNA settings, this means 100,214,929 identical alignment records for the 36.6 million reads with hits. Indexing the Asian elephant genome took 82 s with *galn index* on one GH200, against 2,617 s (44 min) and 3,656 s (61 min) with *bwa index* on AMD EPYC 9755 and AMD EPYC 7742, which is 32× and 45× faster, respectively (**Fig. 1b**). Alignment of the woolly mammoth sequencing reads to the Asian elephant was 6.3× faster using *galn aln* (643 s, 11 min) as compared to *bwa aln* (4,027 s, 67 min) on a full-node comparison within the Arrhenius cluster (**Fig. 1c**). However, speed was drastically lower when performing the same *bwa aln* on the older AMD EPYC 7742 CPU processors on the Dardel cluster (27,285 s, 7.6 hrs).

G-ALN also uses less energy. Aligning the 50 million woolly mammoth reads with aDNA settings took 0.47 kWh on the GPU node, compared with 1.02 kWh on the Arrhenius CPU node and 3.36 kWh on the Dardel CPU node, i.e., a 2.2× and 7.2× lower energy requirement (**Fig. 1d**). The higher power draw of the GPU node (2,567 W vs 892 W on the Arrhenius CPU node) was outweighed by its much shorter run time.

## 4. Discussion

Our results show that G-ALN substantially reduces indexing and alignment times while preserving byte-for-byte compatibility with BWA. The identical outputs across all tested datasets and parameter settings show that the acceleration is achieved without altering the resulting alignment. Indexing a genome can run 32× faster, and subsequent read alignment is 6.3× faster on a 4-year-old NVIDIA GH200 GPU with G-ALN than BWA aln on a 2-year-old 2.7 GHz AMD EPYC 9755 CPU. Since NVIDIA’s latest Vera Rubin GPU architecture delivers up to 4× faster dense compute performance than the GH200, we anticipate that the performance of G-ALN will improve further on newer GPU architectures. The performance gain of G-ALN will therefore evolve over time, much like the unit of measurement on which its pronounced name is derived. Finally, our benchmarking analyses show that G-ALN uses ~2× and ~7× less energy compared to the CPU-based analyses run with BWA aln on the AMD EPYC 9755 and AMD EPYC 7742 processors, respectively, demonstrating a reduced cost and environmental impact of using G-ALN as compared to BWA.

We highlight that G-ALN can easily be incorporated into existing aDNA-processing pipelines, due to its producing identical output files to BWA, and will therefore be included in the next release of the DNAharvester pipeline (Sharif *et al*. 2026). However, G-ALN has potential utility beyond population genomics workflows. The increased speed of both indexing and alignment makes it applicable to large-scale metagenomic workflows, such as those used for sedimentary ancient DNA (sedaDNA), where these steps are key bottlenecks. For such workflows, including the Holi (Pedersen *et al*. 2016) and metaJAM (Johnson *et al*. 2026) pipelines, the Bowtie2 aligner (Langmead and Salzberg 2012) is currently often used due to its superior speed and because, unlike BWA, it can retain read alignments to multiple genomes using the -k parameter. In a future release, we intend to add an equivalent feature to G-ALN so that it can be incorporated into large-scale metagenomics workflows. However, this potential application remains to be evaluated, as our benchmark involved a single mammalian reference genome rather than a large, multi-taxon database typically used in metagenomics. More broadly, GPU acceleration has been, but could be further, extended to other computationally demanding steps in metagenomic pipelines (e.g., GPMeta (Wang *et al*. 2023), MetaCache-gpu (Kobus *et al*. 2021)). Lastly, although G-ALN currently runs only on NVIDIA GPUs, we aim to add AMD support in future releases.

## Code and data availability

The G-ALN source code is publicly available on GitHub (https://github.com/bilalbioinfo/galn). The subsampled, trimmed, and merged sequencing read datasets used for benchmarking are archived at Zenodo: https://doi.org/10.5281/zenodo.23455049. Since G-ALN is adapted from the BWA source code, we recommend that users also cite the original BWA publication (Li and Durbin 2009).

## Author contributions

Conceptualization: BS, KH, TvdV, PDH, LD; Data curation: BS; Formal analysis: BS; Investigation: BS; Methodology: BS, KH; Project administration: LD, BS; Resources: BS, LD; Software: BS; Validation: BS; Visualization: BS; Supervision: LD; Funding acquisition: LD; Writing – original draft: BS, PDH, TvdV, LD; Writing – review & editing: BS, KH, TvdV, PDH, LD.

## Acknowledgements

Computational resources were provided by the National Academic Infrastructure for Supercomputing in Sweden (NAISS), funded by the Swedish Research Council (Arrhenius: NAISS 2026/4-1757; Dardel: NAISS 2026/1-30). The authors also acknowledge additional computational support from GenomeVault AB. Code development was assisted by Claude Opus 5.5 (Anthropic). All code was validated, and correctness was verified by byte-for-byte comparison with stock BWA output.

