## Supplementary Text for "G-ALN: a GPU-accelerated BWA-aln algorithm for palaeogenomics"

**Table S1:** MD5 checksums of the SAI files generated by BWA aln 0.7.19 (CPU node) and G-ALN 0.2.1 (GPU node; default hybrid mode and *--gpu-only*) for woolly mammoth MD090 and steppe mammoth Adycha reads aligned to mEleMax1, and neanderthal Chagyrskaya 8 reads aligned to GRCh38.p14, with aDNA (*-l 1024 -n 0.01 -o 2*) and default settings.

| Dataset | Reads | Parameters | MD5, bwa aln 0.7.19 (CPU node) | MD5, G-ALN 0.2.1 (GPU node) | MD5, G-ALN 0.2.1 --gpu-only (GPU node) |
|---|---|---|---|---|---|
| Mammoth (MD090) | 1,000,000 | aDNA | 70e3f5d6ddc3c682ef8b45a7b3845a08 | 70e3f5d6ddc3c682ef8b45a7b3845a08 | 70e3f5d6ddc3c682ef8b45a7b3845a08 |
| Mammoth (MD090) | 20,000,000 | aDNA | c032f7a23781cbaa16055db8dca84bf9 | c032f7a23781cbaa16055db8dca84bf9 | c032f7a23781cbaa16055db8dca84bf9 |
| Mammoth (MD090) | 50,000,000 | aDNA | 62f417c66dce184c5b26a8467814aa28 | 62f417c66dce184c5b26a8467814aa28 | 62f417c66dce184c5b26a8467814aa28 |
| Mammoth (MD090) | 1,000,000 | default | 6c9926f740dd9f39e0f5152b3fc90b49 | 6c9926f740dd9f39e0f5152b3fc90b49 | 6c9926f740dd9f39e0f5152b3fc90b49 |
| Mammoth (MD090) | 20,000,000 | default | fb40afbe3c52591f0f3a116076c3ca0e | fb40afbe3c52591f0f3a116076c3ca0e | fb40afbe3c52591f0f3a116076c3ca0e |
| Mammoth (MD090) | 50,000,000 | default | 650a5ac43474907a3837388cdc8870f9 | 650a5ac43474907a3837388cdc8870f9 | 650a5ac43474907a3837388cdc8870f9 |
| Mammoth (Adycha) | 1,000,000 | aDNA | 3f04f07aca6d773d2ae8120bed9afc64 | 3f04f07aca6d773d2ae8120bed9afc64 | 3f04f07aca6d773d2ae8120bed9afc64 |
| Mammoth (Adycha) | 20,000,000 | aDNA | e8073fed93ad326a4250978beb15cb45 | e8073fed93ad326a4250978beb15cb45 | e8073fed93ad326a4250978beb15cb45 |
| Mammoth (Adycha) | 50,000,000 | aDNA | 3fa67324e04a8b82b873b2a0c29e914f | 3fa67324e04a8b82b873b2a0c29e914f | 3fa67324e04a8b82b873b2a0c29e914f |
| Mammoth (Adycha) | 1,000,000 | default | be5ac4547b17325c1d756db581cc7f48 | be5ac4547b17325c1d756db581cc7f48 | be5ac4547b17325c1d756db581cc7f48 |
| Mammoth (Adycha) | 20,000,000 | default | e5a3eb7944cfae5cfb3025faefd19d70 | e5a3eb7944cfae5cfb3025faefd19d70 | e5a3eb7944cfae5cfb3025faefd19d70 |
| Mammoth (Adycha) | 50,000,000 | default | d689439720665256b4572ccec1500d6f | d689439720665256b4572ccec1500d6f | d689439720665256b4572ccec1500d6f |
| Chagyrskaya 8 | 1,000,000 | aDNA | 4f7b083d80cc4c92eecc45839b12fd67 | 4f7b083d80cc4c92eecc45839b12fd67 | 4f7b083d80cc4c92eecc45839b12fd67 |
| Chagyrskaya 8 | 2,000,000 | aDNA | 9ff2e6fd9348509935558708d83876ec | 9ff2e6fd9348509935558708d83876ec | 9ff2e6fd9348509935558708d83876ec |
| Chagyrskaya 8 | 5,000,000 | aDNA | e5536ad91e1d89f4ace04b567cb5704b | e5536ad91e1d89f4ace04b567cb5704b | e5536ad91e1d89f4ace04b567cb5704b |
| Chagyrskaya 8 | 1,000,000 | default | f515f72de6e84fdf1be50673d5f378be | f515f72de6e84fdf1be50673d5f378be | f515f72de6e84fdf1be50673d5f378be |
| Chagyrskaya 8 | 2,000,000 | default | 5eb6c5695fabb2bdd6d6afc95d737b34 | 5eb6c5695fabb2bdd6d6afc95d737b34 | 5eb6c5695fabb2bdd6d6afc95d737b34 |
| Chagyrskaya 8 | 5,000,000 | default | 76fe4e04647ed88fb38cf14597d8b48c | 76fe4e04647ed88fb38cf14597d8b48c | 76fe4e04647ed88fb38cf14597d8b48c |

Supplementary fastq datasets are available at Zenodo:
https://doi.org/10.5281/zenodo.23455049
